# Interspecific variation in reproductive and foraging traits for Norwegian raptors

**DOI:** 10.64898/2026.06.28.734957

**Authors:** Eirik Sandvik Halgunset, Jarad Mellard

## Abstract

Arctic and boreal raptor communities are likely to be affected by borealization and other climate change related processes, providing a challenge for ecologists interested in predicting future community structure. However, by combining community assembly theory and species traits, we may be able to better predict future communities. In this study, we analyzed reproductive traits and interspecific variation within those traits and relate it to diet specialization for 29 raptor species breeding in Norway, together with two skua species and three corvid species used for comparative taxa. We also examined published functional response parameters and used specialization related foraging parameters in an illustrative predator-prey model. Specialist raptors produced larger mean clutch sizes and wider reported clutch size ranges than generalists. Specialists also had a higher proportion of fledged per clutch, but the difference was not statistically significant. These results suggest a relationship between diet specialization and reproductive traits which were also observed within phylogenetic orders. Specialist owls (*Strigiformes*) produced higher clutch sizes with a larger clutch range compared to generalist owls. The same pattern was observed for falcons (*Falconiformes*). No clear difference in reproduction was observed for specialist and generalist hawks, kites and eagles (*Accipitriformes*). Corvids expressed clutch sizes similar to that of specialist raptors while having the lowest proportion of fledged per clutch. Differences in foraging traits between specialists and generalists could be distinguished using functional response curves. In the modeled scenarios, a generalist with lower foraging efficiency could coexist with a resident specialist when it had access to prey unavailable to the specialist. The combined empirical and theoretical findings in this study suggest that diet specialization is associated with reproductive traits and may influence the conditions under which predator species invade and coexist.

## Introduction

In northern ecosystems, climate change is altering terrestrial vegetation, causing the spread of woody plants and inducing reductions or even collapses of lemming population cycles (Ims et al., 2013). Additionally, “borealization” may occur, defined as the advancement of boreal species into Arctic or sub-Arctic ecosystems (Criado et al., 2025). These changes are currently reconstructing ecological communities (Root et al., 2003; Verdonen et al., 2026). Changes in vegetation and keystone herbivore populations can have cascading effects on higher trophic levels such as raptors (Carrete et al., 2009; Millon et al., 2014). However, little is known about how raptors are responding and what future Arctic communities will look like. Trait-based community ecology may help identify which species are likely to persist in or colonize changing environments. (Keddy, 1992; McGill et al., 2006). Thus, knowledge of current species traits is key when predicting future Arctic communities. Here, we examine reproductive and foraging traits among raptors breeding or occurring in Norway.

Raptors can be classified as specialists or generalists according to their diets. *Specialists* often consume a single prey type and show a strong numerical and functional response for their preferred prey (Gilg et al., 2006; Hanski et al., 1991). Thus, the abundance of specialists is often linked to the abundance of their main prey (Andersson & Erlinge, 1977; Gilg et al., 2006). *Generalists* use a wider range of prey types, with the ability to switch to a prey type that is currently more abundant (Andersson & Erlinge, 1977; Hanski et al., 1991). However, diet specialization is highly context dependent, and may vary with the organism studied, data used and traits evaluated (Devictor et al., 2010). Previous studies emphasize that diet specialization affects the reproduction of a species (Korpimäki et al., 2020; Pierotti & Annett, 1987). However, interspecific variation in reproduction as a consequence of diet specialization remains poorly understood across species (Korpimäki et al., 2020).

Variation in reproductive traits can be explained by the degree of fluctuations in food availability (Lack, 1954). Raptors with access to stable food resources can express exceptionally stable breeding populations (Newton, 1979). Raptors specialized on cyclic prey, however, tend to breed in higher density and have higher reproductive output when the preferred prey abundance is high (Newton, 1980). Hence, a reproductive trade-off mechanism is associated with diet specialization. In times when main food availability is scarce, specialists can be less efficient at utilizing different resources than generalists (Terraube et al., 2011). Specialists may therefore show greater variability in reproductive output across nests, populations, or years, depending on the availability of their preferred prey. The long-tailed skua for instance, a small rodent specialist, expresses variation in clutch size and breeding success related to rodent abundance (Andersson, 1976). Reproductive success has also been linked to timing and breeding habitat (Terraube et al., 2012) and migration strategies can explain some variation in reproductive strategies (Grist et al., 2017). Migrating and nomadic birds tend to live faster lifestyles (higher fecundity, mature earlier and have shorter lifespans) compared to resident species (Korpimäki, 1992; Soriano-Redondo et al., 2020).

The combined knowledge of reproductive and foraging traits allows us to better understand the ecology of a species and community assemblages of raptors. We can use community assembly theory based on environmental filters and species traits to predict the presence of a species in a habitat (Diamond, 1975; Haefner, 1981; Keddy, 1992). Successful conditions for invasions can be derived from combining species functional traits and environmental conditions in mathematical models simulating borealization events. Trait-based predator–prey models can also be used to explore how differences in foraging efficiency and prey use might influence colonization and coexistence.

The main goals of the study were to examine relationships within and between reproductive and foraging traits in specialist and generalist raptors. We also attempt to understand if and how diet specialization can be used to make community-based predictions. We predicted that specialists would produce larger mean clutches, have wider reported clutch ranges, and have higher proportion of fledglings to clutch size compared to generalists. We also compared these patterns within taxonomic orders where sample size permitted. As a secondary theoretical analysis, we used functional response parameters in a predator-prey model to explore the prey conditions which would allow a generalist predator to invade a community with a resident specialist.

## Method

In this study, we developed two workflows based on data analysis (figure 1). First, reproductive data were collected from the literature for 29 raptor species, two skua species, and three corvid species. The species were separated into diet specialists or generalists based on literature descriptions. Second, foraging data was used to plot functional responses (FR) for four raptor species and one skua species. Diet categorization was determined by trait parameters describing the FR curves. The FR was visually inspected to confirm diet categorization. Thus, some species categorized as specialists in literature might be categorized as generalist based on the plotted functional response. This highlights that diet specialization is context dependent. The reproductive-trait analysis was the primary empirical component of the study, whereas the functional-response and modeling analyses were used as a conceptual extension.

**Figure 1:**
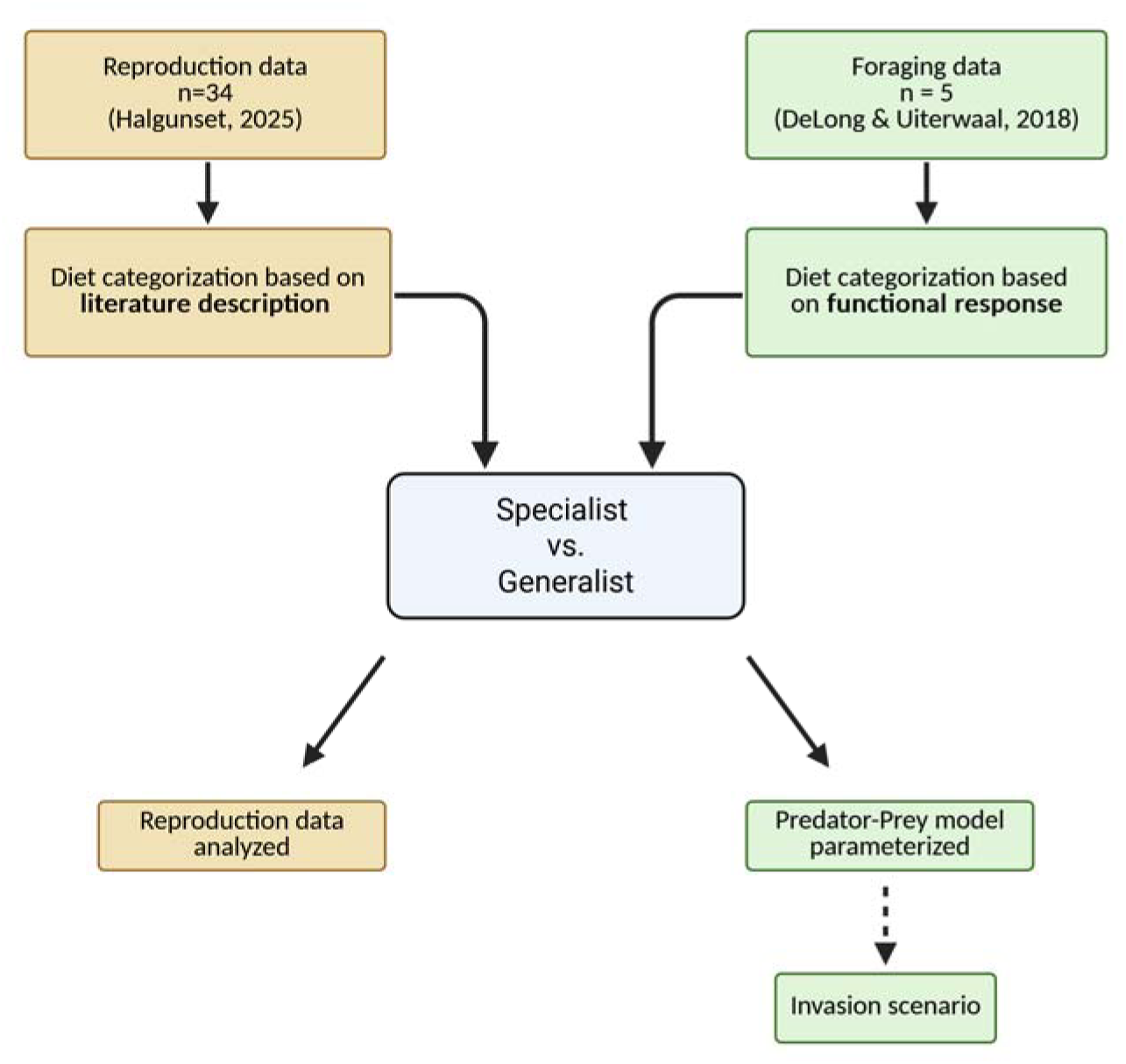
Workflow diagram for this study. On the left, in the yellow boxes, reproduction data for 29 raptor species and two skua species categorized as specialists or generalists based on literature descriptions, were analyzed (in addition to 3 corvid species). To the right, in the green boxes, functional responses for four raptor species and one skua species were plotted, and their diet specialization was determined based on trait data and the functional response. The predator-prey model was parameterized with specialist and generalist specific parameters. Invasion scenarios were simulated with the predator-prey model. Created in https://BioRender.com.

### Study species

Twenty-five raptor species regularly breed in Norway, and four additional species can be considered vagrants or rare breeders (Heggøy & Shimmings, 2020). All 29 raptor species were included in this study in addition to the long-tailed skua, great skua and three corvid species (figure 2). The long-tailed skua was included because it has several ecological similarities to raptors (Andersson, 1976), and a unique adaptation to Arctic tundra, feeding mainly on lemmings during breeding season (Sittler et al., 2011). The great skua has a wider diet and was included to compare to the long-tailed skua. The common raven, hooded crow and the Eurasian magpie were included as examples of omnivore generalists. All species except the Montagùs harrier were listed with migration strategies based on descriptions in Heggøy & Shimmings (2020).

**Figure 2:**
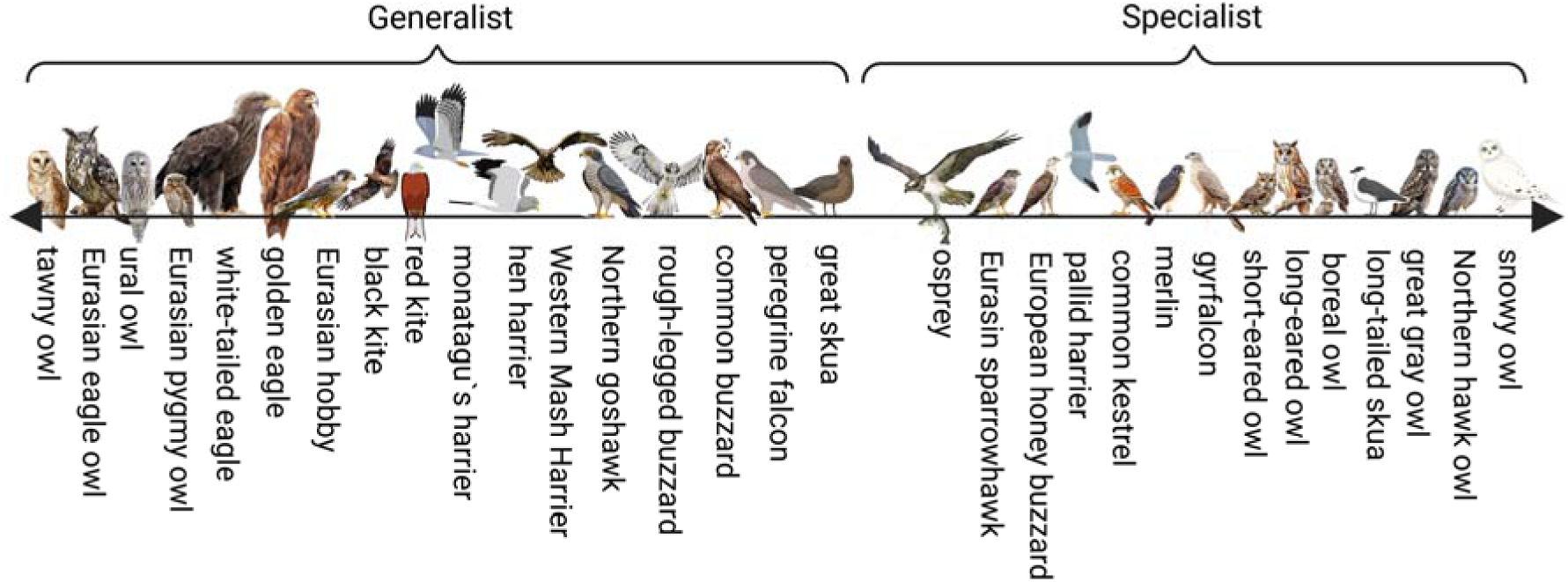
All 29 raptors and two skua species are categorized as generalists or specialists according to their literature descriptions. Placement within each category is for visualization only and does not represent a quantified degree of specialization. Size proportions not to scale. Created in https://BioRender.com.

### Reproductive data

The methods used to identify reproductive-trait data sources and classify species based on literature descriptions are described in Halgunset (2025). A summary is provided here. Clutch size ± standard deviation (SD), number of fledged chicks per clutch ± SD and clutch range were collected from literature references that filled 3 criteria: (1) The data was measured across multiple years. (2) The data included relatively large numbers of nests or clutches, and if several references for one species were available, the reference with the most clutches was chosen. (3) If criteria 1 and 2 were filled, data from studies conducted in Fennoscandia was chosen. We retained one reproductive estimate per species. Hence, the reported standard deviations describe variation within the selected study and do not account for variation among studies or regions.

Proportion fledged was defined as number of fledglings per clutch divided by mean clutch size. Proportion fledged was included because it provides an understanding of the energy invested in reproductive output. The Birds of the World database (Billerman et al., 2025) were used to collect clutch size range for each species. The clutch size range was defined as the maximum and minimum clutch size observed for the species. We acknowledge that reported ranges may depend on sampling effort, hence this variable was interpreted with caution as the published range rather than as a direct estimate of annual variability. For species that did not have clutch size range data available in Birds of the World, the original reference was used. All species in this dataset were categorized as specialists or generalists based on their descriptions in literature (see reference for each species in Halgunset (2025)). It’s important to note is that there exists a range from specialization to generalization within and between different species (Korpimäki et al., 2020), making these classifications difficult and context dependent.

### Foraging Data

FoRAGE (Functional Responses from Around the Globe in all Ecosystems) database: a compilation of functional responses for consumers and parasitoids (Uiterwaal & DeLong, 2021; Uiterwaal et al., 2018) was used to collect foraging data for four raptor species and one skua species. The snowy owl and long-tailed skua were feeding on lemmings. The short-eared owl, long-eared owl and common kestrel were feeding on voles. Type-Ⅱ functional response curves were reconstructed using the reported handling time and attack rate parameters. Handling time was defined as handling time per resource and attack rate as space clearance rate per predator. Other species in the FoRAGE database were not raptors or not wild raptors feeding on wild prey and were thus not included. Species were separated into specialists (n=2) or generalists (n=3) based on the species-specific parameters of handling time and attack rate and plots of functional responses were used to visually confirm this classification. To simplify, we calculated a ratio of generalist/specialist handling time 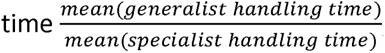, and implemented these ratio-parameters in the predator-prey model.

### The Predator-Prey Model

Population dynamics between predator(s) and prey(s) were simulated using the Rosenzweig-MacArthur predator-prey model (Rosenzweig & MacArthur, 1963). The model was intended as an illustrative and an exploration of specialization-related mechanisms rather than as a species-specific prediction. The model was modified to include two prey species (N_1_ and N_2_) and two predator species (P_1_ and P_2_). A specialization coefficient (β_i_) was added to the model to describe prey preference. The model is given by:

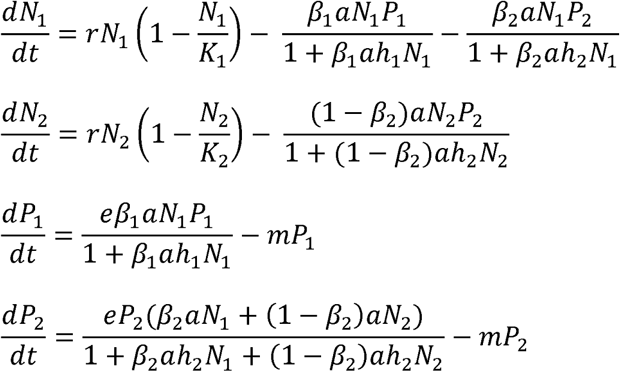

where *r* is intrinsic growth rate, *K* is environmental carrying capacity, *a* is attack rate on prey, *h_i_* is handling time of prey, *e* is the conversion efficiency for food intake to new predators, *m* is the per-capita mortality rate and *β_i_* is the specialization coefficient. We assume that prey N_1_ and N_2_ are of similar body size and therefore consider handling time to be related to the predator and not the prey. We also determined that attack rate parameters are constant for both predators on both prey species but are always multiplied with the specialization coefficient *β_i_* which differs for P_1_ and P_2_.

#### Invasion scenario

To extract conditions for successful invasions, the predator-prey model was simplified with the following assumptions:

i. The growth rate for the invading predator —— ——— needs to be greater than 0 for a successful invasion
ii. Conversion efficiency (e) and mortality rate (m) is equal and constant for all predators and prey
iii. Handling time (h) for the generalist predator is twice as long compared to the specialist predator (calculated from foraging data)
iv. Attack rate (a) was set equal to 0.02 for all predators but was always multiplied with a specialization coefficient β ∈ [0,1), where 1 is a predator entirely specialized on one prey
v. A generalist predator (P_2_) invades a stable predator-prey community with a resident specialist (P_1_) and its prey (N_1_) at their equilibrium

Initial values for the invasion scenario were r=4, N_1_=100, K_1_=1000, a_1_=0.02, h_1_=0.1, e=0.04 and m = 0.15. The values were chosen to give the predator-prey model an equilibrium, simplifying the analysis. Equilibrium was verified numerically when changes in all state variables were negligible over the final fifty timesteps (see figure A3).

### Statistical analysis

The difference between specialists, generalists and corvids was analyzed with the following response variables: clutch size mean, proportion of fledged per clutch size and clutch range difference. The corvids were separated from the raptors in the analysis because we wanted to directly compare corvid reproduction strategies to that of raptor generalists and specialists. To identify differences between specific categories, Tukey’s honest significant difference (HSD) was used. ANOVA was used to test for difference in clutch size means and proportion fledged per clutch between specialist, generalists and corvids. The number of species and observations within taxonomic groups is limited, hence we did not conduct a formal phylogenetic comparative analysis. However, we repeated comparisons within orders where sample sizes permitted. We consider these order-specific analyses as exploratory because they do not fully account for phylogenetic non-independence. Within *Strigiformes* one influential value was observed. Within *Falconiformes* and *Accipitriformes* homoscedasticity was violated and influential values were observed. Assumptions were not fully met for Falconiformes and Accipitriformes, thus results for these orders were interpreted cautiously. No test was performed on *Charadriiformes* because only two species were present. Additional Anova tests were performed on the clutch range with clutch range difference as response variable. No statistical analyses were performed on the difference in reproduction between migration strategies because many strategies had one or few observations.

The data was analyzed using R version 4.4.2 (R Core Team, 2024). The R script is found in the supplementary material.

## Results

### Reproduction Traits

The median species-level mean clutch size was 6.0 eggs for corvids, 3.0 eggs for generalists, and 4.8 eggs for specialists. Mean clutch size differed among corvids, generalists, and specialists (F_2,31_ = 7.21, p = 0.003). Levenès test for unequal variance of clutch size mean was not statistically significant (Levenès test: F_2,31_ = 1.75, p = 0.19). Specialists had larger mean clutch sizes than generalists (p = 0.02). Corvids had larger mean clutch sizes than generalists (p = 0.01). The estimated difference in mean clutch size between specialists and generalists was 0.2–2.5 eggs based on the 95% confidence interval (Appendix Figure A1).

Specialists had the highest proportion fledged, but the difference between specialists and generalists was not statistically significant (95% CI: -0.06, 0.16, p = 0.50). Proportion fledged differed among the three categories (F_2,24_ = 7.84, p = 0.002). Corvids had lower proportion of fledged per clutch than specialists (95% CI: 0.12, 0.55, p = 0.001) and generalists (95% CI: 0.05, 0.48, p = 0.01).

Figure 5 shows the clutch sizes and clutch ranges for all 34 species included in the study. The size differed between specialists and generalist owls (*Strigiformes*) (F_1,8_ = 6.47, p = 0.034). The mean clutch size for generalist owls was 3.98 eggs (t-value = 7.29, SE = 0.54, p < 0.001), while specialist owls lay, on average, 1.77 more eggs compared to generalist owls (t-value = 2.54, SE = 0.70, p = 0.03). The difference in mean clutch sizes between generalists and specialists within *Falconiformes* (F_1,3_ = 4.48, p = 0.124) *and Accipitriformes* (F_1,12_ = 0.85, p = 0.374) was not statistically significant. Because some model assumptions were not met for *Falconiformes* and *Accipitriformes*, these results should be interpreted cautiously.

The clutch range was significantly different between specialist and generalist *Strigiformes (*F_1,8_ = 8.65, p = 0.02) and *Falconiformes (*F_1,3_ = 45.34, p = 0.006) but not within *Accipitriformes* (F_1,12_ = 3.30, p = 0.09). The long-tailed skua and the great skua had the narrowest clutch size range with 1-2 eggs, followed by honey buzzard, peregrine falcon and white-tailed eagle with a range of 1-3 eggs.

### Functional response

The reconstructed functional response curves were similar for the common kestrel, long-eared owl and short-eared owl. The snowy owl and long-tailed skua, however, had higher consumption rates and approached their asymptotic consumption rates at lower prey densities than the other species. The ratio for handling time between generalist and specialist was calculated as: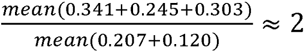 (see Uiterwaal & DeLong (2021) for data). In this five-species dataset, the median handling time of species classified as relative generalists was approximately twice that of species classified as relative specialists. The long-tailed skua had a lower attack rate and lower handling time, saturating at higher consumption rates with increasing prey densities compared to the snowy owl. Based on their functional-response parameters and curves, the snowy owl and the long-tailed skua were classified as relative specialists. In contrast to their literature-based descriptions in the reproductive trait analysis, the common kestrel, short-eared owl and long-eared owl were classified as relative generalists based on the functional response.

### Invasion scenario example

Three scenarios were simulated (Figure 3 and Appendix figure A3-A6), and their main results are summarized below. The resident specialist predator and its prey reached equilibrium after several timesteps. Equilibrium values from scenario Ⅰ were used in scenario Ⅱ and Ⅲ. Under the selected parameter values, the resident specialist excluded the generalist when both predators competed for a single prey species, because the specialist’s shorter handling time allowed growth at lower attack rates (Appendix figure A4 and A5). In scenario Ⅲ the generalist was able to switch between two prey types. When the generalist had access to an alternative prey species unavailable to the specialist, it achieved growth, even though the handling time was twice as long as the specialist. Under the parameter values given in the figure caption, the simulation resulted in coexistence between the generalist and the specialist (appendix figure A6).

**Figure 3:**
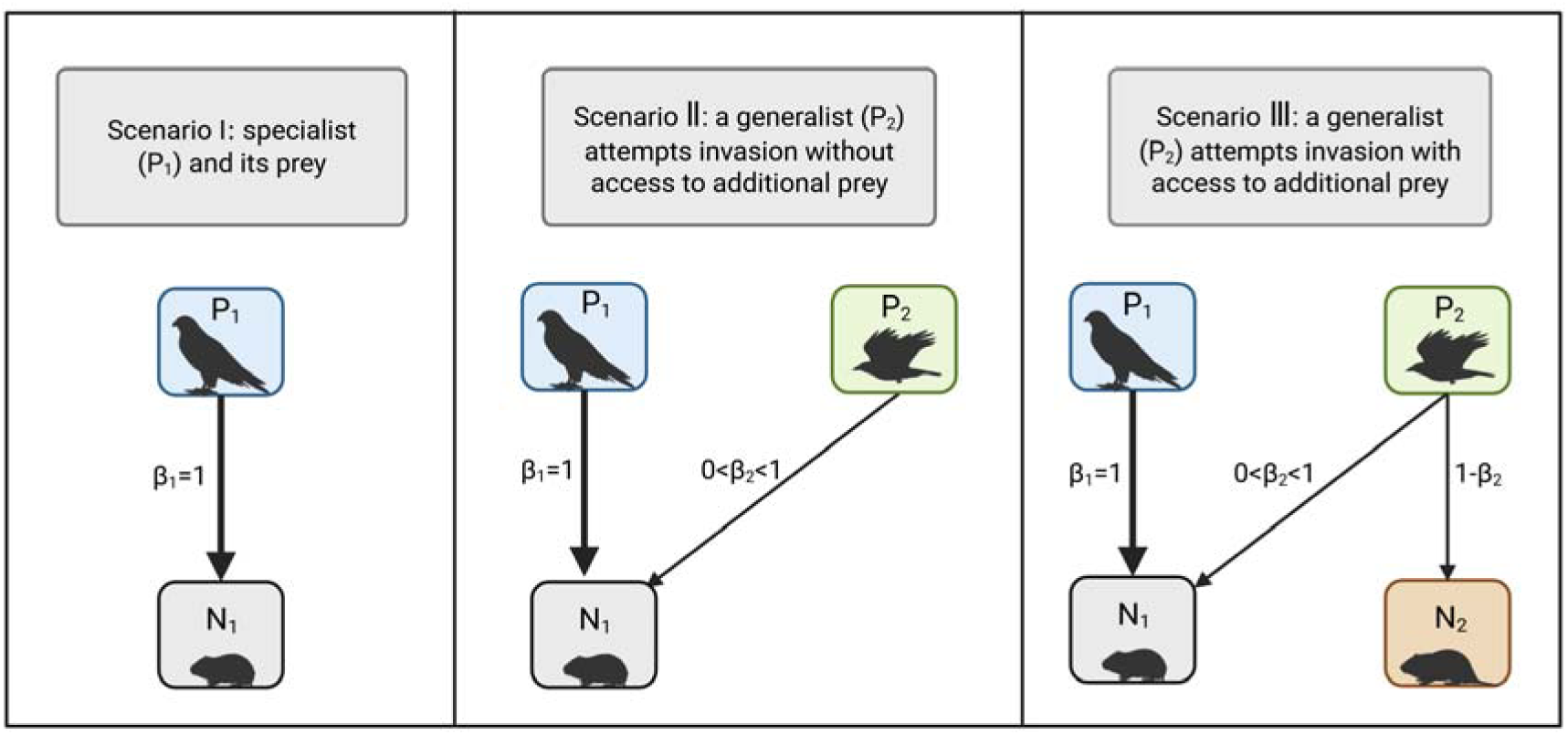
Three different scenarios were simulated. A specialist with a single prey (Ⅰ). A resident specialist competing with a generalist attempting invasion with access to one prey species (Ⅱ). A resident specialist competing with a generalist attempting invasion with access to two prey species (Ⅲ).

**Figure 4:**
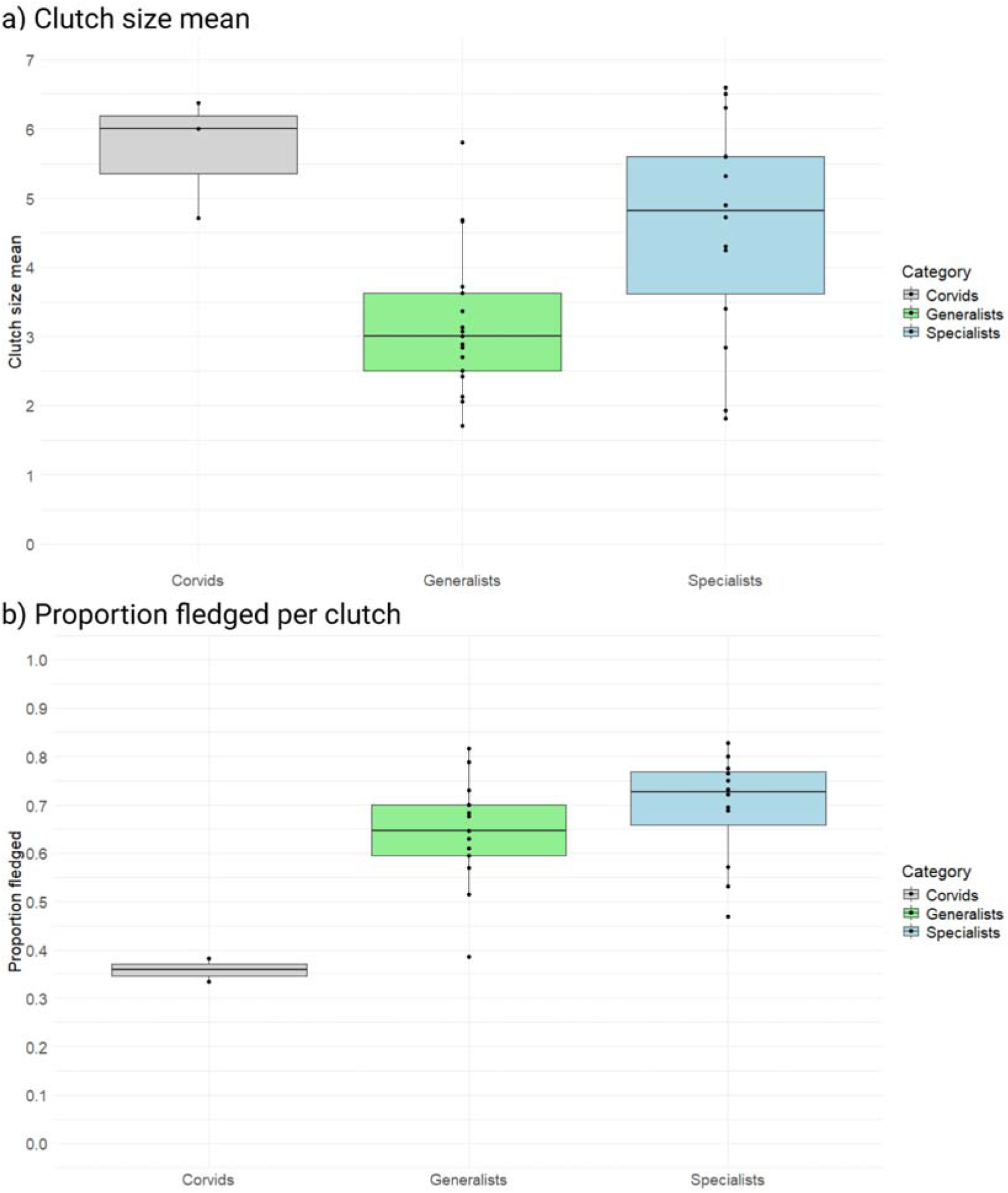
Reproductive traits by diet category. a) Species-level mean clutch size for specialists: n = 14, generalists: n =17, and corvids n=3. The pygmy owl was identified as an outliner. b) shows the proportion of fledged chicks per clutch for each category. All species were included except: Eurasian pygmy owl, Northern hawk-owl, peregrine falcon, hooded crow and the long-tailed skua due to missing data in reference. The short-eared owl and long-eared owl were identified as outliners.

**Figure 5:**
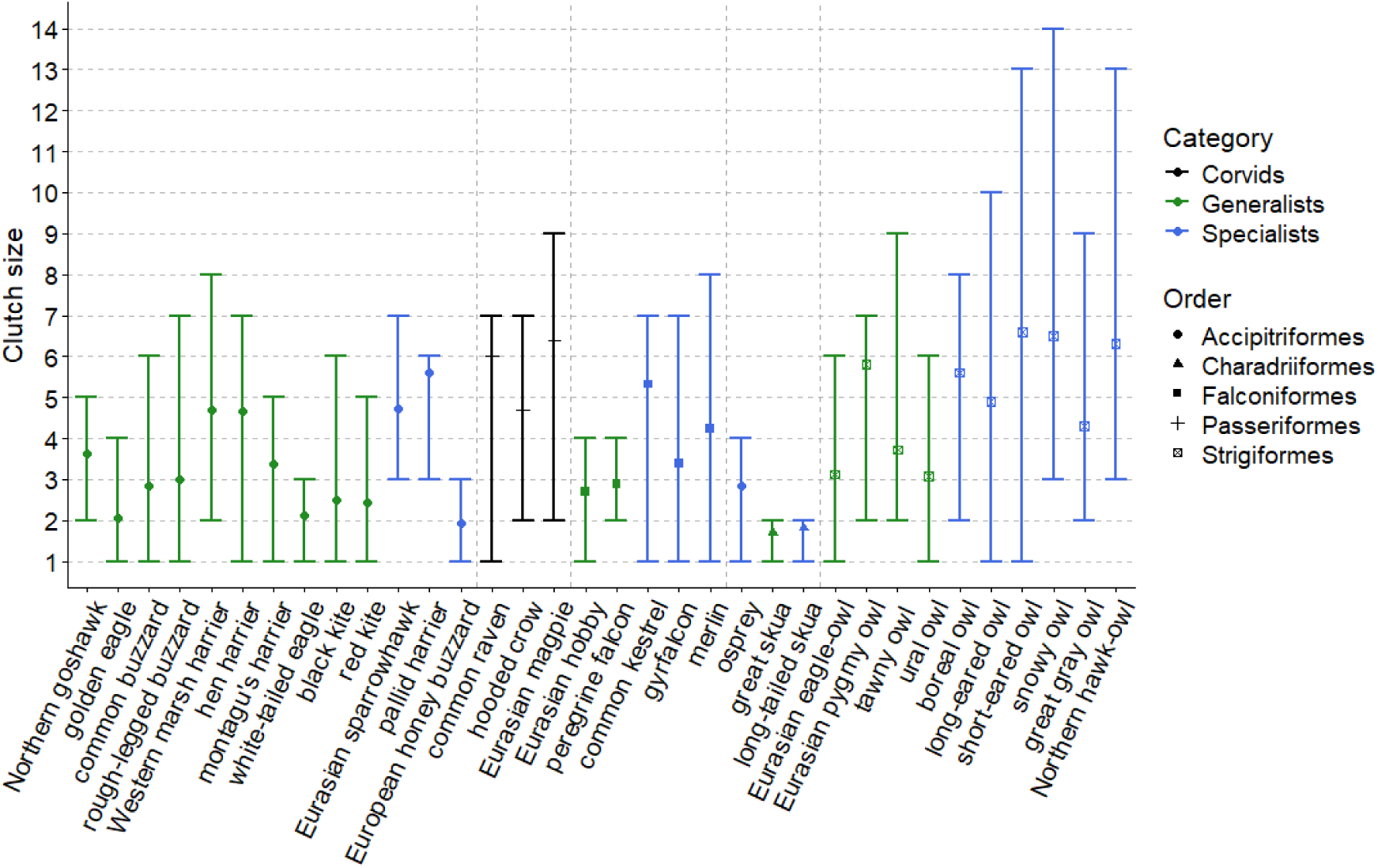
Clutch size range and species-level mean for all species included in the study. The y-axis represents the minimum and maximum clutch size reported in Birds of the World or, when unavailable there, in the selected primary reference. The species are divided into diet categories specialists, generalists and corvids, represented by color. The symbols represent the mean clutch size per species listed in phylogenetic orders.

**Figure 6:**
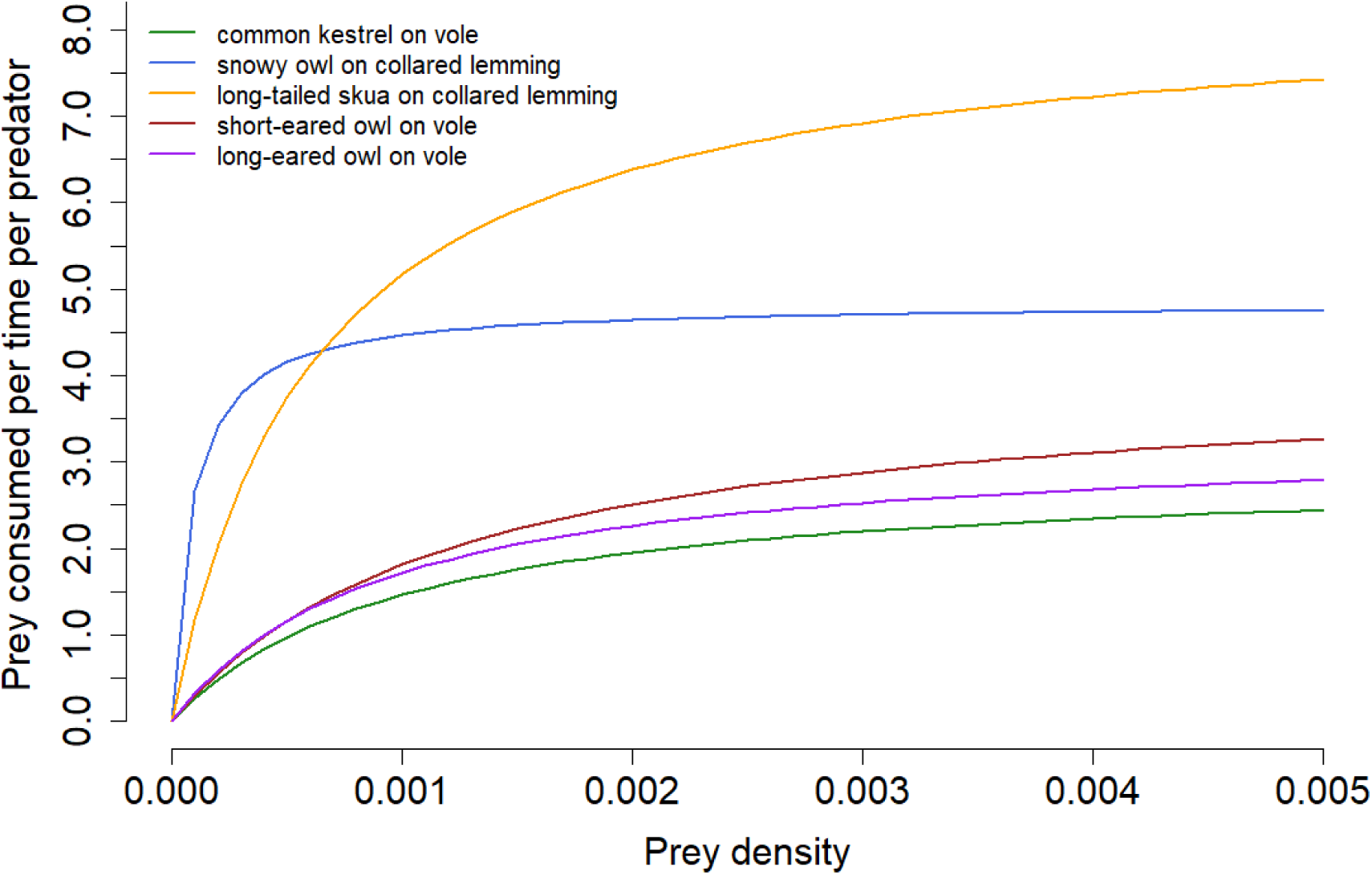
Holling type II functional-response curves reconstructed for four raptor species and one skua species using median attack rates and handling times from the FoRAGE database. Curves show estimated consumption rate as a function of prey density; consumption approaches an asymptote as prey density increases.

**Figure 7:**
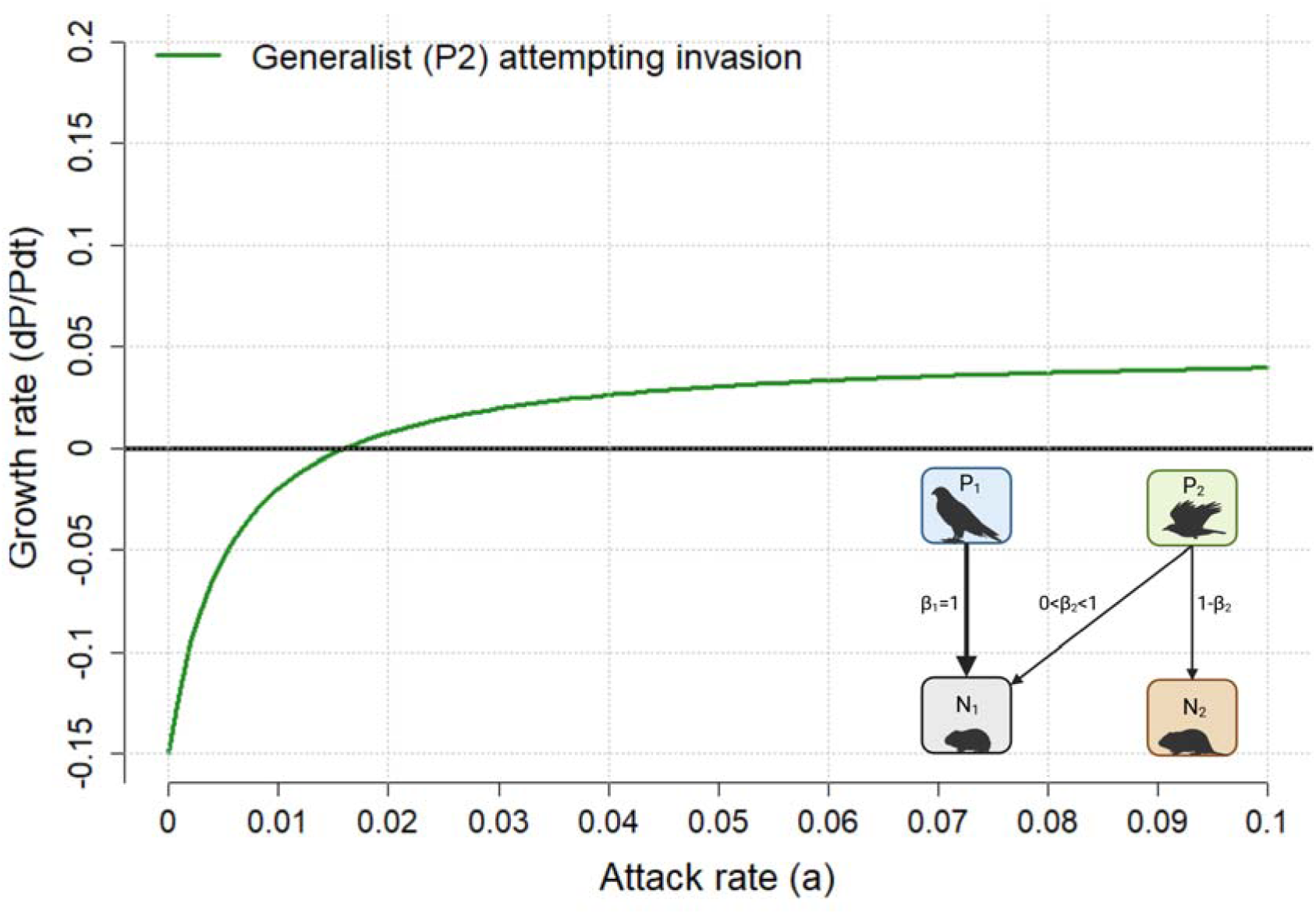
Per-capita growth rate (dP/Pdt) of the generalist predator plotted across attack rates ranging from 0 to 0.1 when two prey species were available. The diet specialization coefficient β describes allocation toward prey N1. A low β means that a predator is more specialized on N2 because β=1 is a predator entirely specialized on N1. Parameter values in the model were N1=300, N2=1000, e=0.04, m = 0.15, while the handling time (time per resource) was 0.2 for the generalist (twice that of the specialist. Extracted from the foraging data) and β2=0.1.

In this simulation, the generalist’s establishment depended on access to alternative prey and on the effective attack rate, which was modified by the specialization coefficient. Under the selected parameter values, the generalist did not coexist with the specialist when only one prey species was available. In this scenario, the persistence of the generalist depended on access to an alternative food source unavailable to the specialist. The attack rate is influenced by the degree of specialization on the prey. Thus, a specialization coefficient 0 < β_2_ < 0.5 means that the generalist achieved an attack rate that gave population growth. Establishment was favored when the generalist directed more of its foraging effort toward the alternative prey than toward the specialist’s preferred prey.

## Discussion

In this study, we have analyzed the interspecific variation in reproductive and foraging traits as a consequence of diet specialization for raptors breeding/occurring in Norway. In line with our predictions, specialists produced significantly higher clutch sizes, had wider reported clutch ranges and tended towards higher proportion of fledged per clutch compared to generalists. Previous research with fewer species is consistent with our findings (Korpimäki, 1992; but see Korpimäki et al., 2020 for contradiction; Pierotti & Annett, 1987; Terraube et al., 2011). The difference in reproduction between specialist and generalist raptors was interpreted across 5 phylogenetic orders. However, no phylogenetic corrections were made in the analysis, and results should therefore be interpreted with caution. Additionally, foraging traits for four raptors and one skua species were evaluated and implemented into a predator-prey model. We showed that a successful invasion is determined by a species’ ability to switch to prey unavailable to the native species, which is determined by its diet specialization and foraging traits.

Specialist raptors expressed more variation in reproduction compared to generalist raptors. Within phylogenetic orders, specialist owls (*Strigiformes)* produced higher clutches than generalist owls. Specialists owls generally have shorter llifespans and are smaller bodied (Korpimäki, 1986). Such owl species produce, on average, greater clutch sizes compared to larger species (Korpimäki et al., 2020). Similar results were shown within *Falconiformes*. Specialist falcons produced larger clutches and expressed wider clutch ranges compared to the generalists. Within *Accipitriformes,* no clear differences in reproduction were observed, although a general trend of more variation for specialists was present. However, the number of *Accipitriformes* species was skewed towards generalists. These interspecific trait variations could be explained by a reproductive trade-off mechanism associated with diet specialization. Generalists produce smaller clutches, fledge on average a lower proportion of chicks while expressing minimal variation due to their ability to switch to alternative prey when main prey abundance is low. Specialists, however, fledge a higher proportion of chicks, but are only able to do so when main prey are abundant (Korpimäki et al., 2020). It’s important to note that diet specialization is context dependent. For example, the boreal owl is considered a resident generalist in central Europe. In northern Fennoscandia, however, the boreal owl behaves like a nomadic specialist (Korpimäki, 1986). It’s also important to note that these results should be interpreted with caution because no phylogenetic corrections were implemented.

Nomadic specialist owls produced larger clutch sizes compared to migrating specialists as well as resident generalists. The snowy owl, a nomadic specialist, expressed clutch ranges between 3-14 eggs. Anderson (1980) suggested that nomads are better suited to fluctuating prey populations because of their ability to adjust clutch sizes (Andersson, 1980). The long-tailed skua is a migrant species that behaves like a specialist raptor. However, it only lays 1-2 eggs per clutch. Egg experiments with the long-tailed skua revealed that skuas are in fact able to produce at least 4 eggs per clutch, but adding 1 egg to their clutches resulted in hatching failure (Andersson, 1976). This suggests evolutionary constraints on skua reproduction. First of all, the incubation capacity hypothesis is a viable explanation for clutch sizes in the skua family (*Stercorariidae*) (Reid, 1987). Skuas might be unable to brood more than 2 eggs per clutch due to their brooding method, incubating one egg between one foot and a brood patch (Andersson, 1976). Furthermore, 1-2 eggs per clutch for skuas could be an adaptation to egg-predation rates (Godfray et al., 1991). Finally, above average clutch sizes yield lower reproductive output because of insufficient parental resource foraging (Williams, 1966). The small clutch sizes of skuas could also be explained as an adaptation to minimize energy loss if prey abundance is low.

A clear difference in reproductive traits was observed between the corvids and the specialist/generalist raptors. In this study, the raven, hooded crow and magpie were included. These species can be considered omnivore generalists with an opportunistic lifestyle adapted to uncertain food supplies (see Daw et al., 2025). However, the clutch size of corvids was similar to that of specialist raptors. Despite the large clutch sizes, corvids had the lowest proportion of fledged per clutch of all species included in this study. Some possible explanations for this might come from the brood reduction hypothesis (Lack, 1947, 1954) and insurance hypothesis (Forbes, 1990). Corvids express hatching asynchrony (Holyoak, 1967), which may be adaptive for individuals unable to predict resource levels in their upcoming nestling season (Pijanowski, 1992). Hatching asynchrony, brood reduction, and insurance eggs may explain the high clutch sizes and low fledgling number of corvids. The reproductive trade-off mechanism associated with diet specialization could explain how some extreme generalists, such as many corvids, produce large clutches as insurance for poor and unpredictable conditions.

We used a predator-prey model, parameterized with foraging data, to conduct simulations of invasions. Specialist/generalist foraging traits were evaluated based on the properties of the functional response (FR), which to our knowledge, has not been done before. This method revealed that lemming specialists had higher consumption rates than species feeding on voles. In previous research, predator-prey models with a type Ⅰ FR have been simulated (Rivera-Estay et al., 2024). Here we implemented a Type-Ⅱ FR, contributing to a more realistic predator invasion description. We showed that the successful invasion of a generalist was determined by 2 conditions: the ability to switch to alternate prey and the attack rate multiplied with a specialization coefficient which represents the degree of specialization on alternate prey. These invasion conditions give rise to several implications. First, two species using the same resource the same way cannot coexist (Hardin, 1960). Thus, the presence of an invading generalist could lead to competitive exclusion of native predators (Rivera-Estay et al., 2024), as shown in scenario Ⅱ. However, previous simulations show that specialists are only able to outperform generalists when the specialist is N times more effective catching their preferred prey (N being the number of preys in the system) (Araujo & Moura, 2022). In other words, a specialist must be twice as good at catching a preferred prey compared to a generalist who has access to two preys. Thus, coexistence between a generalist and a specialist occurs under restricted conditions (Araujo & Moura, 2022).

We used the invasion analyses to simulate a borealization-related invasion event. In northern ecosystems, the increase of generalist predators is driven by climate change and borealization (Ims et al., 2019). There seems to be contradictions and confusion in the literature as to which species, or rather which traits, allow species to successfully invade, especially in birds. Marino & Bellard (2023) found that invasive bird species had larger clutch sizes and tended to be dietary generalists compared to native species. Sol et al. (2012), however, argued that brood size is a better predictor to estimate invasion success. Species with low brood size had higher invasion success with the argument that such species may prioritize future rather than current reproduction (Sol et al., 2012). Both arguments can be explained by reproductive trade-offs associated with diet specialization. Corvids, for example, had large clutch sizes, low proportion of fledged per clutch and expressed low variation. Ravens have increased their abundance and distribution in northern ecosystems (Harju et al., 2021), which may be explained by their generalist diet and opportunistic lifestyle.

In conclusion, we propose that the reproductive trade-off mechanism associated with diet specialization can be extended to include invasion success. Thus, community-based predictions can be made from traits linked to diet specialization. Specialists produce larger clutches, have a higher proportion of fledged per clutch while expression more year-to-year variation due to fluctuations of main prey abundance. Specialists may thus fledge a large number of chicks only in favorable conditions. Such species are less likely to successfully invade. Generalists, however, produce smaller clutches, have a lower proportion of fledged per clutch while expressing less year-to-year variation due to their ability to switch to alternate prey. Generalists may thus be more likely to invade because of their ability to utilize alternate prey and consistently breed. However, we have also showed that some extreme diet generalists, like many corvids, produce large clutch sizes while fledging few chicks. We suggest that invasion success can also favor species that are able to consistently fledge chicks even if the brood size is small. However, we did not analyze invasion simulations with fluctuating prey, but rather stable prey populations which can be viewed as a peak rodent year.

Future research should focus explicitly on categorizing more species based on quantitative trait data or functional responses as diet specialists/generalists to further examine trade-off mechanisms on reproduction and invasion success. Phylogenetic correction could be implemented into the analysis to account for evolutionary relationships, but this requires data. A sensitivity analysis on parameters could also be implemented to view how each parameter affects the outcome in simulations. Furthermore, detailed data to parameterize predator-prey models are needed. In this study, the conversion efficiency of prey to new predators was assumed to be constant. The reproduction output could be implemented into this coefficient. Such coefficient could represent how much energy the predator puts into producing a clutch of eggs and reproductive output (fledgling probability). In a simplified framework an organism can be viewed as input/output systems with foraging tactics as the input and reproduction tactics as the output (Pianka, 1976). Furthermore, a multispecies functional response curve (Abrams, 2022) for generalists consuming a variety of prey types should be analyzed using data which we currently lack. Finally, future research should examine how a generalist invasion would affect fluctuating specialist predator-prey cycles, as Boreal generalists are increasing their abundance in the Arctic due to climate change.

## Supporting information

R-script and readme file

## Appendix

**Figure A1.**
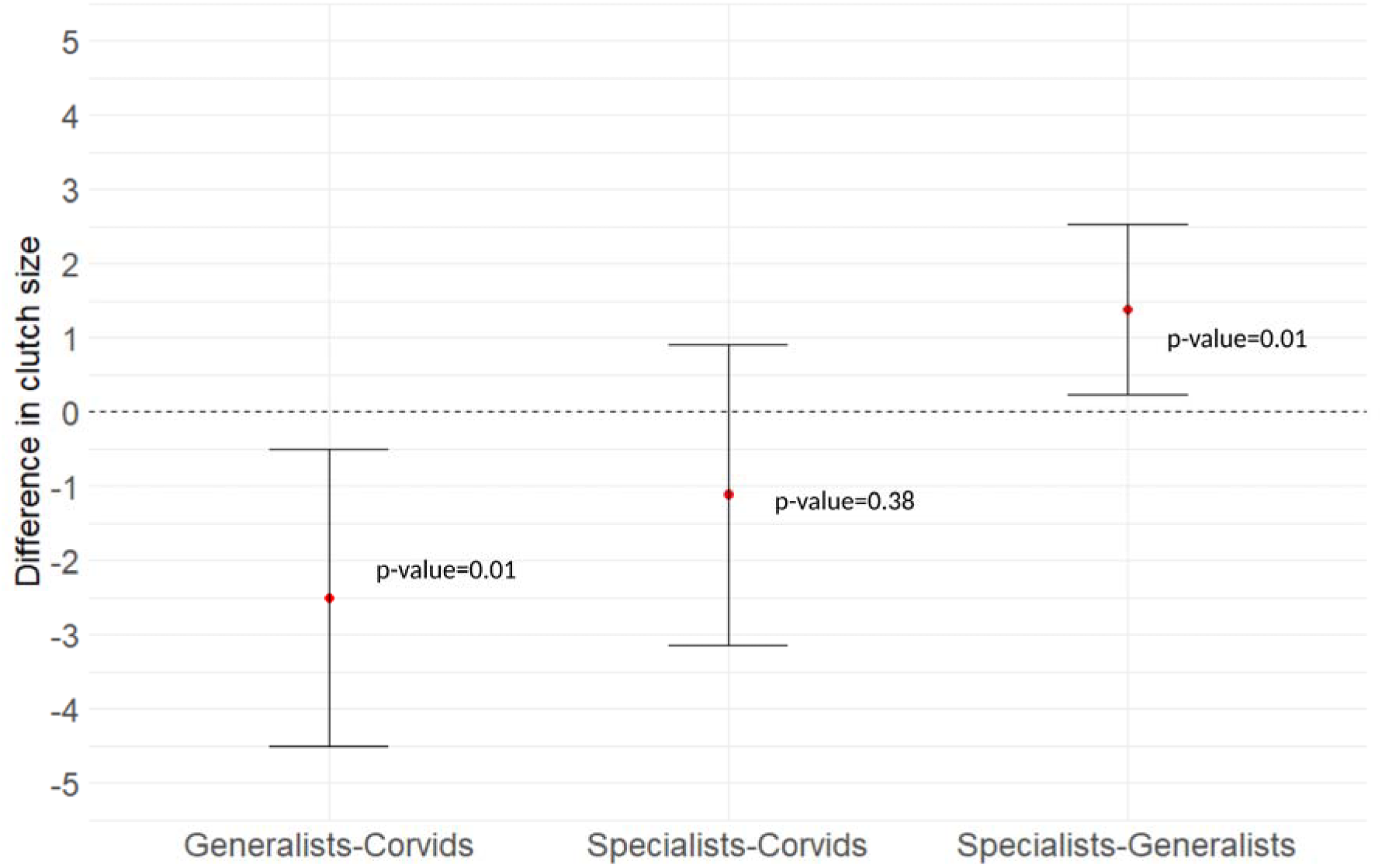
: 95% confidence interval from anova test on difference in clutch size between corvids, generalists and specialists.

**Figure A2.**
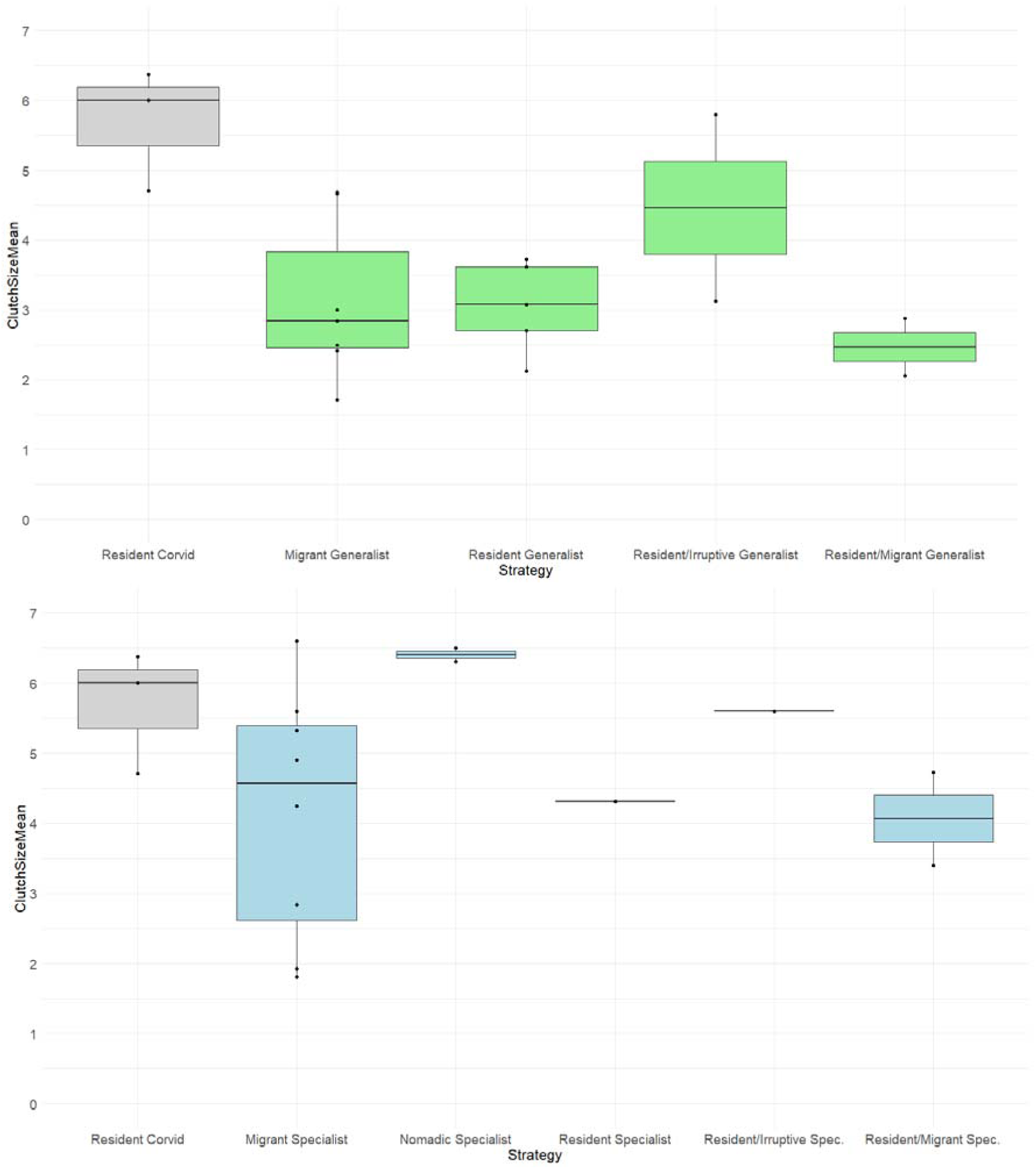
: Species divided into specialists (n=14), generalists (n=17) and corvids (n=3) in addition to migration strategies. No statistical differences between diet categories and migration strategies can be accounted for due to few observations and model assumption violation although differences can be observed. Nomadic specialists produce higher clutch sizes compared to migrants and resident specialists. For generalists, no clear difference is observed between the migration strategies.

**Figure A3.**
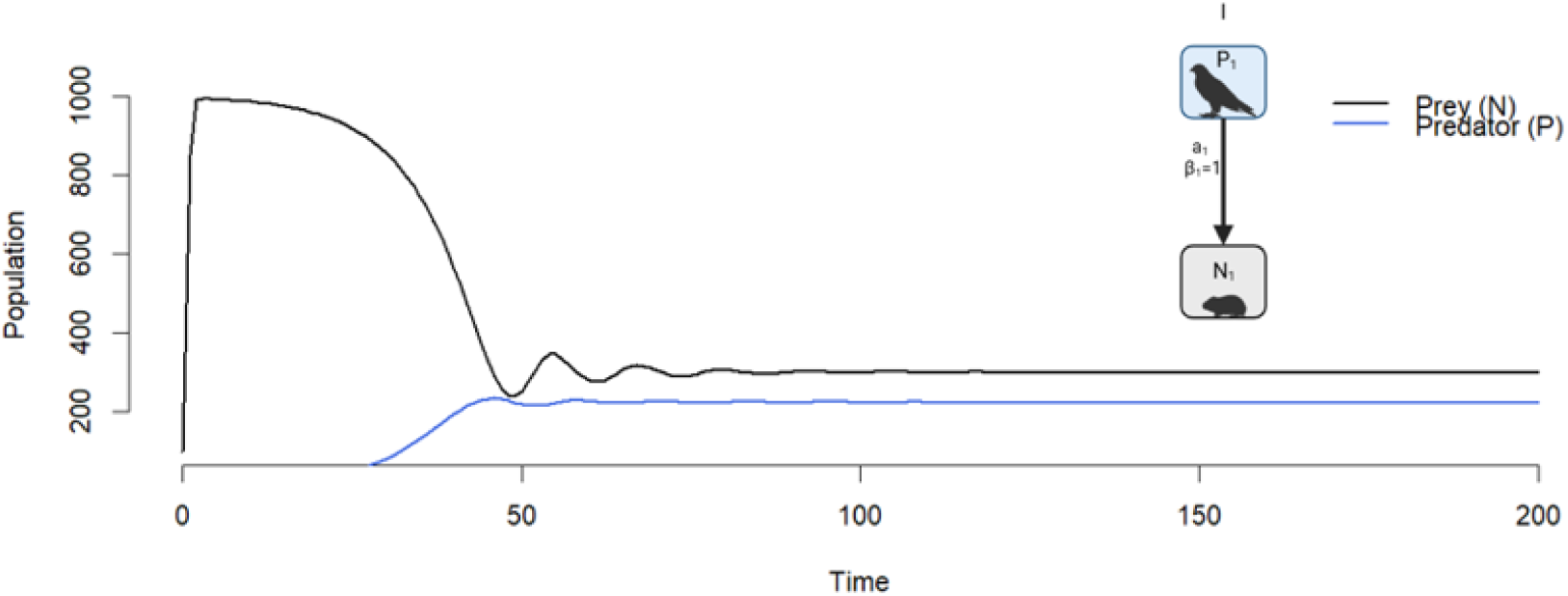
: Simulation of the first scenario: a specialist predator and its prey. The y-axis represents the predator and prey population, and the x-axis shows the number of time steps. Prey parameter values: r=4, N=100, K=1000. Predator parameters: P=10, e=0.04, β=1, a=0.02, h=0.1, m=0.15. Figure A3 shows a simple simulation of the first scenario: a single specialist and its prey. The model reaches equilibrium after several timesteps. These equilibrium values were used in the second scenario.

**Figure A4.**
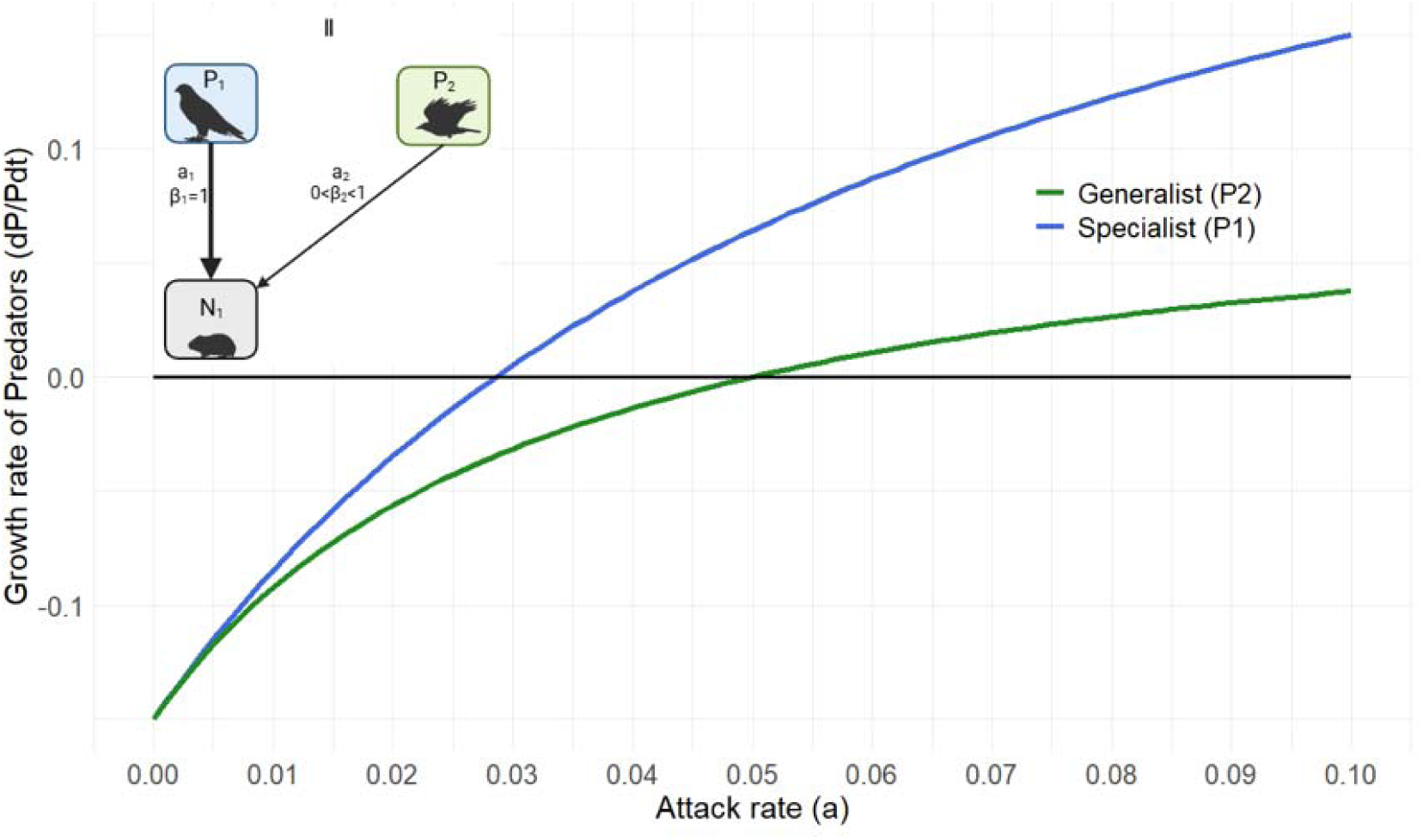
: The growth rate (dP/Pdt) for a specialist and generalist predator with attack rates from 0 to 0.10. Equal parameter values for P_1_ and P_2_ were used: N=300, e=0.04, m = 0.15. For the handling time (h), the generalist had a handling time twice as long compared to a specialist. It was assumed that the only parameter differentiating between the specialist and generalist was the handling time and attack rate times β. Here, β was not included in the model because only one prey was available. When competing for single prey, the specialist could achieve population growth with lower attack rates compared to the generalist (figure A4). The specialist could have an attack rate of 0.02 and still achieve population growth. The generalist, however, needed to have a minimum attack rate of 0.05 to achieve population growth. When competing for a single prey, the generalist needed an attack rate more than 2 times faster than the specialist to achieve population growth. Data from the FoRAGE database shows that the specialists had an attack rate 4 times higher than the generalist, while the handling time was 2 times faster.

**Figure A5.**
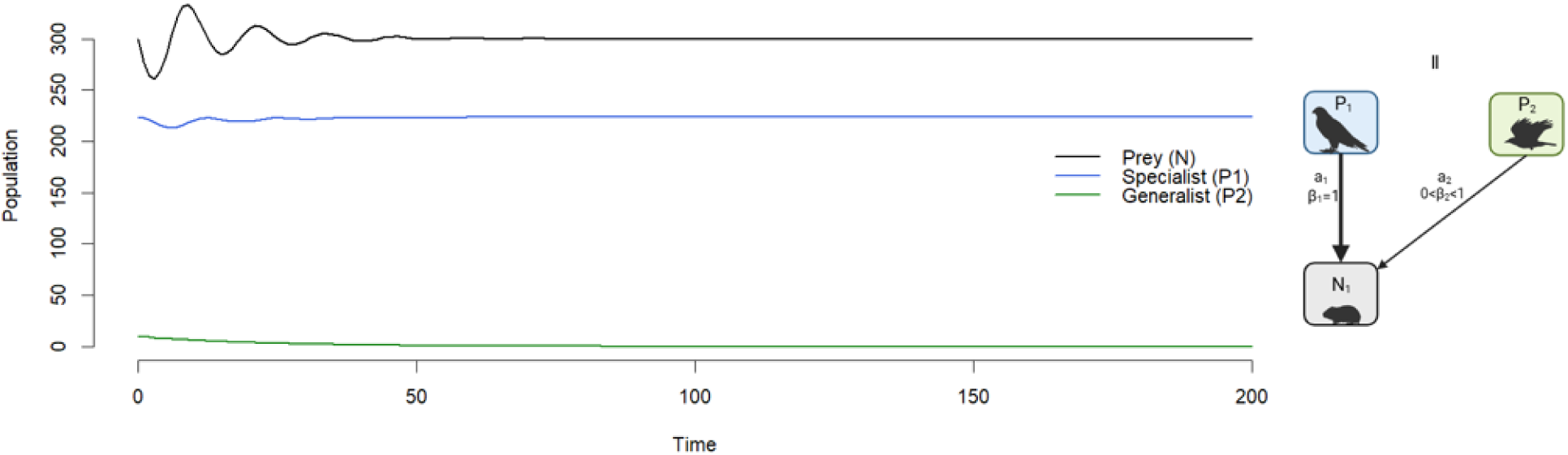
: Simulation of the second scenario: a specialist predator and its prey with a second generalist predator added. The y-axis represents the predator and prey population, and the x-axis shows the number of time steps. Prey parameter values: r=4, N=300, K=1000. Specialist parameter values e=0.04, β_1_=1, a=0.02, h_1_=0.1, m = 0.15. Generalist parameter values: e = 0.04, β_2_=0.9, h_2_=0.1*2, m = 0.15. In the second scenario (figure A5), the specialist predator outcompeted the generalist predator. As the growth rate of P_2_ never exceeded 0, the invading population of generalist predator declined, leading to extinction. The native specialist predator reached the same equilibrium presented in scenario one.

**Figure A6.**
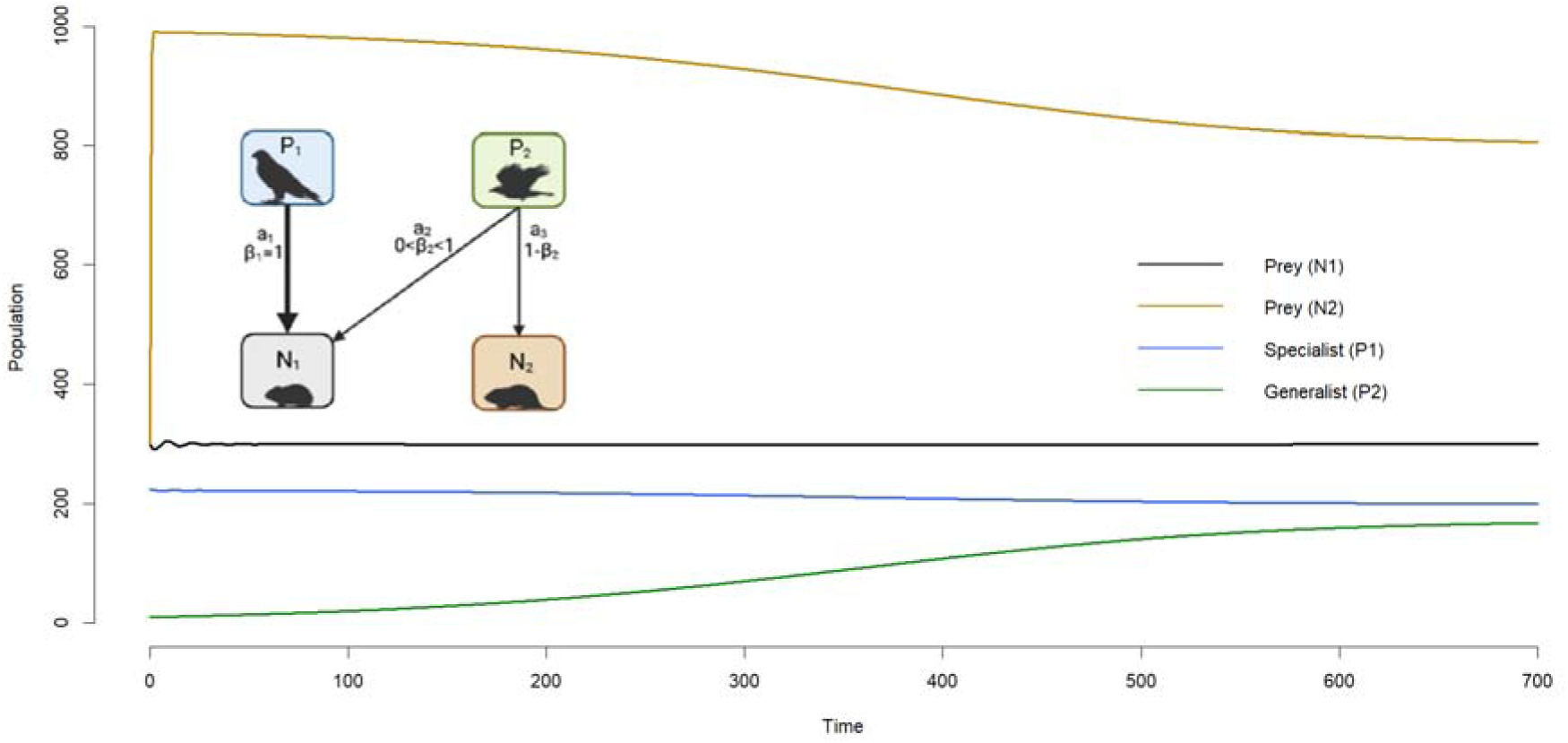
: Simulation of a generalist invading a specialist predator feeding on one prey type. The y-axis represents the predator and prey populations, and the x-axis shows the number of time steps. Prey parameter values: r=4, N_1_=300, N_2_=300, K_1_=K_2_=1000. Specialist parameter values e=0.04, β_1_=1, a=0.02, h_1_=0.1, m = 0.15. Generalist parameter values: e = 0.04, β_2_=0.1, h_2_=0.1*2 m = 0.15. Simulation of a generalist invading a specialist resulted in co-existence between both predators and their prey. The specialization coefficient β_2_ were set to 0.1. The specialist predator only preyed on its preferred prey (N_1_) and had a specialization coefficient β_1_ = 1. In the simulation, the attack rate of both predators was set equal, and the only parameter differentiating the specialist and generalist was the handling time and specialization coefficient.

## Notes

https://doi.org/10.18710/NJEDNQ

### Competing Interest Statement

The authors have declared no competing interest.

### Summary of Updates

Removed the front page with contact information; fixed one reference

