## Supplementary material for "Interspecific variation in reproductive and foraging traits for Norwegian raptors": R-script and readme file: r_script.R

```
# =============================================================================
# Title:        Data analysis for reproductive and foraging traits for raptors 
#               breeding in Norway
# Paper:        Interspecific variation in reproductive and foraging traits for
#               Norwegian raptors
# Authors:      Eirik Sandvik Halgunset & Jarad Mellard 
# Affiliation:  Univeristy of Tromso
# Date:         2026-09-23
# R version:    4.4.2
# Description:  This script contains all code to reproduce analysis and results 
#               regarding the reproduction and foraging traits for raptors 
#               breeding in Norway as well as invasion scenarios. 
# =============================================================================

library(deSolve)
library(ggplot2)
library(forcats)
library(car)
library(dplyr)

# ==============================================================================
# === Read the reproduction data================================================

# Read the data from DataverseNo, doi.org/10.18710/NJEDNQ
raptor <- read.csv("reproduction_data_raptors.csv", sep = ";", dec = ",")
raptor <- raptor[1:34,] #Filters out empty columns

# === Boxplots Specialists, Generalists and Corvids=============================

# Clutch size mean
ggplot(raptor, aes(x = Category, y = ClutchSizeMean, fill = Category))+
  geom_boxplot()+
  geom_point()+
  theme_minimal()+
  theme(axis.title.x = element_blank())+
  labs(y = "Clutch size mean")+
  scale_fill_manual(values = c("Generalists" = "lightgreen", "Corvids" = "lightgrey", 
                               "Specialists" = "lightblue")) +
  scale_y_continuous(limits = c(0,7), breaks = seq(0,7,by=1))+
  theme(
    legend.text = element_text(size = 14),
    axis.text.x = element_text(size = 14),      
    axis.text.y = element_text(size = 14),        
    axis.title.y = element_text(size = 16),
    title = element_text(size = 16))

# Proportion fledged 
ggplot(raptor, aes(x = Category, y = ProportionFledged, fill = Category))+
  geom_boxplot()+
  geom_point()+
  theme_minimal()+
  theme(axis.title.x = element_blank())+
  labs(y = "Proportion fledged")+
  scale_fill_manual(values = c("Generalists" = "lightgreen", "Corvids" = "lightgrey", 
                               "Specialists" = "lightblue")) +
  scale_y_continuous(limits = c(0,1), breaks = seq(0,1,by=0.1))+
  theme(
    legend.text = element_text(size = 14),
    axis.text.x = element_text(size = 14),      
    axis.text.y = element_text(size = 14),        
    axis.title.y = element_text(size = 16),
    title = element_text(size = 16))

# === Clutch range for each species ============================================

# Clutch range for each species 
ggplot(raptor, aes(x = fct_reorder(CommonName, interaction(Family, Category, Genus), 
                                   .fun = as.character), ymin = ClutchRangeMin, 
                   ymax = ClutchRangeMax, color = Category)) +
  geom_errorbar(width = .7 , lwd = 0.9) +
  geom_point(aes(y = ClutchSizeMean, shape = Order), size = 2)+
  geom_vline(xintercept = c(13.5, 16.5, 21.5, 24.5), 
             linetype = "dashed", color = "darkgray")+
  theme_classic()+
  labs (y = "Clutch size",
        shape = "Order",)+
  theme(axis.text.x = element_text(angle = 65, vjust = 1, size = 10, hjust = 1), 
        axis.title.x = element_blank())+
  scale_color_manual(values = c("Generalists" = "forestgreen", 
                                "Corvids" = "black", 
                                "Specialists" = "royalblue"))+
  scale_y_continuous(breaks = seq(0, 15, by = 1))+
  theme(
    legend.text = element_text(size = 14),
    axis.text.x = element_text(size = 15),      
    axis.text.y = element_text(size = 14),    
    axis.title.y = element_text(size = 16),
    title = element_text(size = 16))+
  theme(panel.grid.major.y = element_line(color = "gray", linetype = "dashed"))


# ==============================================================================
# === Clutch size per category =================================================

ggplot(raptor, aes(x = Order, y = ClutchSizeMean, fill = Category))+
  geom_boxplot()+
  theme_minimal()+
  theme(axis.title.x = element_blank())+
  labs(y = "Clutch size mean")+
  scale_fill_manual(values = c("Generalists" = "lightgreen", 
                               "Corvids" = "lightgrey", 
                               "Specialists" = "lightblue")) +
  scale_y_continuous(limits = c(0,7), breaks = seq(0,7,by=1))+
  theme(
    legend.text = element_text(size = 14),
    axis.text.x = element_text(size = 14),      
    axis.text.y = element_text(size = 14),        
    axis.title.y = element_text(size = 16),
    title = element_text(size = 16))

# ==============================================================================
# === Statistical analysis of reproduction (Categories) ========================
# Clutch size 
lm_mod <- lm(ClutchSizeMean~Category, data=raptor)
par(mfrow = c(2,2))
plot(lm_mod) # evaluate assumptions 

leveneTest(ClutchSizeMean~Category, data=raptor) # test for homoscedasticity

# Use a one-way ANOVA
ANOVA_clutch <- aov(ClutchSizeMean~Category, data=raptor)
summary(ANOVA_clutch)
# Tukey Honest significant difference to check which categories differ
tukey_result <- TukeyHSD(ANOVA_clutch, conf.level = 0.95)

# Proportion fledged 
lm_mod1 <- lm(ProportionFledged~Category, data=raptor)
par(mfrow = c(2,2))
plot(lm_mod1) #Evaluate assumetions 

leveneTest(ProportionFledged~Category, data=raptor) #Test for homoscedasticity

# Use a one-way ANOVA
ANOVA_fled <- aov(ProportionFledged~Category, data=raptor)
summary(ANOVA_fled)
TukeyHSD(ANOVA_fled)

# === Plotting 95% confidence intercal for clutch size =========================
# Plot the 95 CIs for clutch size 
tukey_df <- as.data.frame(tukey_result$Category)

# Add comparison names as a column
tukey_df$Comparison <- rownames(tukey_df)

# Plotting the intervals
ggplot(tukey_df, aes(x = Comparison, y = diff))+
  geom_point(color = "red", size = 2)+
  geom_hline(yintercept = 0, linetype = "dashed")+
  geom_errorbar(aes(ymax = upr, ymin = lwr), width = 0.3)+
  theme_minimal()+
  scale_y_continuous(limits = c(-5,5), breaks = seq(-5,5, by = 1))+
  labs (y = "Difference in clutch size", x = "")+
  theme(
    axis.text.x = element_text(size = 16),      
    axis.text.y = element_text(size = 16),    
    axis.title.y = element_text(size = 16))
# ==============================================================================
# === Statistical analysis of reproduction (Phylogenetic orders) ===============

# Separate the data into phylogenetic orders
owl <- raptor %>%
  filter(Order == "Strigiformes")
falc <- raptor %>%
  filter(Order == "Falconiformes")
hawk <- raptor %>%
  filter(Order == "Accipitriformes")

# Linear models and anova for clutch size ~ order
# Owls
lm1 <- lm(ClutchSizeMean~Category, data = owl)
plot(lm1)
anova(lm1)
summary(lm1)

# Falcons 
lm2 <- lm(ClutchSizeMean~Category, data = falc)
plot(lm2)
anova(lm2)
summary(lm2)

# Hawks, eagles and kites 
lm3 <- lm(ClutchSizeMean~Category, data = hawk)
plot(lm3)
anova(lm3)
summary(lm3)

# Linear models and anova for clutch range ~ order
# Owls
lm1.1 <- lm(ClutchRangeDiff~Category, data = owl)
plot(lm1.1)
anova(lm1.1)
summary(lm1.1)

# Falcons 
lm2.1 <- lm(ClutchRangeDiff~Category, data = falc)
plot(lm2.1)
anova(lm2.1)
summary(lm2.1)

# Hawks, eagles and kites
lm3.1 <- lm(ClutchRangeDiff~Category, data = hawk)
plot(lm3.1)
anova(lm3.1)
summary(lm3.1)
# ==============================================================================

# === Functional Response ======================================================
par(mfrow = c(1,1))

# Data fitted are collected from the FoRAGE database doi:10.5063/F1RX99KB 
# Values fitted are attack rate and handling time (fitted a, and fitted h (day)) 
# in the database

func_res <- function(a,h,N){
  R <- a*N/(1+(a*h*N))
return(R)
}

# Fitted values
N = seq(0,0.005,by=0.0001)
forage <- data.frame(
  kestrel <- func_res(a =2920,h =  0.341, N = N),
  snowy <- func_res(a=59400 ,h =0.207  , N = N),
  skua <- func_res(a=13600 ,h =0.120  , N = N),
  short<- func_res(a = 3260 ,h = 0.245 , N = N),
  long <- func_res(a = 3590,h = 0.303 , N = N),
  prey_den <- seq(0,0.005,by= 0.0001 )
)

# Plotting the functional response
plot(forage$prey_den, forage$kestrel, type = "l", col = "forestgreen", lwd = 2,
     xlab = "Prey density", ylab = "Prey consumed per time per predator", 
     ylim = c(0, 8), axes = F,
     cex.lab = 1.5)
# Add lines for other species
axis(2,seq(0,9,by = 0.5), cex.axis = 1.5)
axis(1, cex.axis = 1.5)
lines(forage$prey_den, forage$snowy, col = "royalblue", lwd = 2)
lines(forage$prey_den, forage$skua, col = "orange", lwd = 2)
lines(forage$prey_den, forage$short, col = "brown", lwd = 2)
lines(forage$prey_den, forage$long, col = "purple", lwd = 2)
# Add a legend
legend("topleft", legend = c("common kestrel on vole", 
                             "snowy owl on collared lemming", 
                             "long-tailed skua on collared lemming", 
                             "short-eared owl on vole", "long-eared owl on vole"),
       col = c("forestgreen", "royalblue", "orange", "brown", "purple"),
       lwd = 2, bty = "n")

# ==============================================================================

# === Invasion models ==========================================================

# P is predator abundance
# a is attack rate on the prey
# h is handling time of the prey
# e is the energetic conversion of prey consumed to predators
# m is the predator mortality rate

# Growth rate function
test <- function(e,b,a,N1,N2,h,m){
  rdP <- e*(b*a*N1 + (1-b)*a*N2)/(1+(b*a*h*N1) + (1-b)*a*h*N2) - m 
  return(rdP)
}

# A data frame with growth rates across different attack rates for each predator. 

df <- data.frame(
  
  rdP_Gen = test(e=0.04, b = 0.1,a = seq(0, 0.1, by = 0.001), 
               N1=300,N2 = 1000, h = 0.1*2, m = 0.15),
  
  rdP_Spe = test(e=0.04, b = 1,a = seq(0, 0.1, by = 0.001), 
                 N1=300,N2 = 1000, h = 0.1, m = 0.15),
  
  attack_rate = seq(0,0.1,by = 0.001))

# Each growth rate plotted against different attack rates 
plot(df$attack_rate, df$rdP_Gen, type = "l", col = "forestgreen", lwd = 3,
     xlab = "Attack rate (a)", ylab = "Growth rate (dP/Pdt)", 
     xlim = c(0, 0.1), ylim = c(-0.15, 0.2), axes = F, cex.lab = 1.6)
abline(h = 0, col = "black", lwd= 3)
axis(1, at = seq(0, 0.1, by = 0.01), labels = seq(0, 0.1, by = 0.01), cex.axis = 1.4)
axis(2, at = seq(-0.2, 0.2, by = 0.05), labels = seq(-0.2, 0.2, by = 0.05), cex.axis = 1.4)
grid(nx = NULL, ny = NULL, col = "gray", lty = "dotted", lwd = 0.8)
legend("topleft", legend = c("Generalist (P2) attempting invasion"), 
       col = c("forestgreen"), lwd = 3, cex = 1.5,  bty = "n")


# === 1 predator, 1 prey model==================================================
# Creating the predator-prey model
RM_pred <- function(time, y, p){
  N <- y[1]; P <- y[2]
  with(as.list(p),{
    dN <-  r*N*(1-N/K) - a*N*P/(1+a*h*N)
    dP <- e*a*N*P/(1+a*h*N) - m*P
    return(list(c(dN, dP)))
  })
}

# Initial parameters
t <- 0:200; y0 <-c(N=100, P=10)
p <- list(r=4, K=1000, a=0.02, e=0.04, m=0.15, h=0.1) 
pred1 <- as.data.frame(ode(y0, t, RM_pred, p))

plot(pred1$time, pred1$N, type = "l", xlab = "Time", ylab = "Population", 
     lwd = 2, bty = "n", ylim =c(0,1000))
lines(pred1$time, pred1$P, col = "royalblue", lwd = 2)

legend("topright", legend = c("Prey (N)", "Predator (P)"), 
       col = c("black", "royalblue"), lwd = 2, bty = "n")

# === 2 predators, 1 prey model=================================================
# Adding P2 to the model
RM_pred <- function(time, y, p){
  N <- y[1]; 
  P1 <- y[2];
  P2 <- y[3]
  with(as.list(p),{
    dN <- r*N*(1-N/K)-a1*N*P1/(1+a1*h1*N)-a2*N*P2/(1+a2*h2*N) 
    dP1 <- e1*a1*N*P1/(1+a1*h1*N) - m1*P1
    dP2 <- e2*a2*N*P2/(1+a2*h2*N) - m2*P2
    return(list(c(dN, dP1,dP2)))
  })
}

t <- 0:200; y0 <-c(N=300, P1=224, P2 = 10)
p <- list(r=4, K=1000,
          a1=0.02, e1=0.04, m1=0.15, h1=0.1, #P1
          a2=0.02*0.9, e2=0.04, m2=0.15, h2=0.1*2) #P2, attack rate multiplied with specialization coefficient = 0.9

out <- as.data.frame(ode(y0, t, RM_pred, p ))

plot(out$time, out$N, type = "l", col = "black", lwd = 2, bty = "n", 
     ylim = c(0, max(out$N, out$P1, out$P2)),
     xlab = "Time", ylab = "Population")
lines(out$time, out$P1, col = "royalblue", lwd = 2)
lines(out$time, out$P2, col = "forestgreen", lwd = 2)
legend("right", legend = c("Prey (N)", "Specialist (P1)", "Generalist (P2)"),
       col = c("black", "royalblue", "forestgreen"), lwd = 2, bty ="n")

# === 2 predator, 2 prey model==================================================
# Adding N2 to the model
RM_pred <- function(time, y, p){
  N1 <- y[1]; 
  N2 <- y[2];
  P1 <- y[3];
  P2 <- y[4]
  with(as.list(p),{
    dN1 <- r*N1*(1-N1/K1)-a*N1*P1/(1+a*h1*N1)-b*a*N1*P2/(1+b*a*h2*N1) 
    dN2 <- r1*N2*(1-N2/K2)-(1-b)*a*N2*P2/(1+(1-b)*a*h2*N2) 
    dP1 <- e*a*N1*P1/(1+a*h1*N1) - m*P1 
    dP2 <- (e*(b*a*N1*P2 + (1-b)*a*N2*P2))/(1+b*a*h2*N1 + (1-b)*a*h2*N2) - m*P2 
    return(list(c(dN1, dN2, dP1,dP2)))
  })
}
# Defining the parameters in the model
# Initial values
t <- 0:700; y0 <-c(N1=300, N2 = 300,  P1=224, P2 = 10)
p <- list(r=4, r1= 4, K1=1000, K2 = 1000,
          a=.02, e=0.04, m=.15, h1=0.1, h2=0.1*2, b = 0.1) 

# Solving the differential equation with deSolve package
out <- as.data.frame(ode(y0, t, RM_pred, p))

# Plotting invasion scenario 3
plot(out$time, out$N1, type = "l", col = "black", lwd = 2.5, 
     ylim = c(0, max(out$N1, out$N2, out$P1, out$P2)),
     xlab = "Time", ylab = "Population", 
     bty = "n")
lines(out$time, out$P1, col = "royalblue", lwd = 2.5)
lines(out$time, out$P2, col = "forestgreen", lwd = 2.5)
lines(out$time, out$N2, col = "darkgoldenrod", lwd = 2.5)
legend("right", legend = c("Prey (N1)", "Prey (N2)", "Specialist (P1)", 
                           "Generalist (P2)" ),
       col = c("black", "darkgoldenrod", "royalblue", "forestgreen"), 
       lwd = 2, bty = "n")

# ==============================================================================
```
