## Supplementary material for "Interspecific variation in reproductive and foraging traits for Norwegian raptors": R-script and readme file: README.txt

This readme file was generated on 2026-06-27 by Eirik Sandvik Halgunset

GENERAL INFORMATION


Author
Name: Eirik Sandvik Halgunset 
ORCID: 0009-0003-8908-7818
Institution: UiT The Arctic University of Norway
 

Co-investigator 
Name: Jarad Mellard
ORCID:0000-0003-1824-919X
Institution: UiT The Arctic University of Norway


DATA & FILE OVERVIEW

File List: r_script.R

DATA-SPECIFIC INFORMATION FOR: r_script.R

The R script contains code to reproduce analysis and figures for "Interspecific variation in reproductive and foraging traits for Norwegian raptors"

The code was written under R version 4.4.2 using the following packages: deSolve, ggplot2, forcats, car and dplyr.

The reproduction data can be downloaded from: https://doi.org/10.18710/NJEDNQ
